# Applying an automated NMR-based metabolomic workflow to unveil strawberry molecular mechanisms in vernalization

**DOI:** 10.1101/2024.02.13.580094

**Authors:** Andrea Fernández-Veloso, Jaime Hiniesta-Valero, Alejandra Guerra-Castellano, Laura Tomás, Miguel A. De la Rosa, Irene Díaz-Moreno

## Abstract

Metabolomics is the discipline that aims to determine the whole metabolic profile of a complex mixture. These studies are useful to capture the physiological status of an organism at a given moment. Even though the main technique used in metabolomics is Mass Spectrometry coupled to chromatography, in recent years, interest in Nuclear Magnetic Resonance (NMR) is increasing because of some benefits of NMR, i.e., it is a non-invasive, highly reproducible, and inherently quantitative technique. However, difficulties in data analysis comprise one of the main reasons that hinder the standardization of NMR for metabolomic analysis in research. In this work, we applied an automated workflow for NMR-based metabolomic analysis for the study of vernalization in strawberry. Vernalization is a key process in obtaining a successful strawberry crop, however, the molecular mechanisms behind it remain still unknown. We expect this work to improve the knowledge of crop metabolism —specifically the vernalization process— while promoting the use of NMR in conjunction with computational tools for agriculture studies.

## 1. Introduction

Metabolomics has emerged as a powerful scientific discipline in the study of small metabolites present in an organism or biological system. Unlike genomics and proteomics, which focus on genes and proteins respectively, metabolomics focuses on the downstream products of genomic, transcriptomic, and proteomic processes (Johnson *et al*. 2018). These small chemical compounds, such as amino acids, lipids, and secondary metabolites, provide an accurate picture of the physiological state of an organism at a given time (Gavaghan *et al*. 2000).

Nuclear Magnetic Resonance (NMR) spectroscopy, routinely used to determine protein structure, is emerging as a useful technique, suitable for metabolite detection in complex mixtures. This technology has been employed mainly in biomedicine to obtain an image of the metabolic status of the organism through spectrum profiles. However, the intricacy of these spectrum profiles obtained in this field, as well as the link between the profile and biological interpretation, continue to be major analytical challenges, complicating its establishment as a routine methodology (Gavaghan *et al*. 2000).

Despite the availability of other analytical techniques such as Mass Spectrometry (MS), gas chromatography, and high or ultrahigh-performance liquid chromatography, NMR is gaining popularity due to its various benefits (Emwas *et al*. 2019). NMR has associated major advantages such as being non-invasive, not requiring separation of compounds from biological mixtures, high reproducibility, inherently quantitative nature, and relatively low cost (García-Villaescusa *et al*. 2018; Abreu *et al*. 2020). In parallel, computational advances have been and continue to be developed to meet the requirements for rapid automatic analysis of the increasingly complex spectra arising from metabolomics studies (Moco, 2022). For all of the above, the use of NMR for metabolomic analysis is becoming increasingly common, mostly in food science and biomedicine (Wu *et al*. 2022; Chowdhury *et al*. 2023). In this work, we propose its usage in a novel application field, for the analysis of plant material from strawberry.

Strawberry (*Fragaria × ananassa* Duch) is an herbaceous, perennial, and stoloniferous plant, which has great climatic adaptability, showing its best potential, especially in warm areas. Its fruit has a high economic value in the world market due to its organoleptic properties and the wide variety of uses that can be given to it (Bahukhandi *et al*. 2022). However, its cultivation is not always easy because it relies on a set of environmental factors, which, besides, may differ depending on the variety used. Among these factors are the photoperiod, cold exposure time during dormancy, the temperature during vegetative development and flowering, and the type of soil. All of these will influence the crop yield and the quality of the harvested product (Morales *et al*. 2017).

One of the parameters that has been seen to be extremely important in obtaining successful crops is the cumulative required number of hours under 7 °C during dormancy, a term known as vernalization. There are several studies demonstrating that it is beneficial for the plant to go through such process (Oviedo *et al*. 2020), since it triggers a change in its physiology, preparing it for flowering and fruit production, and thus promoting good development and abundant fruiting (Costa *et al*. 2014). However, the molecular and metabolic mechanisms that comprise vernalization, which may be the key to improving current agricultural practices and crops, are not fully understood.

It should be noted that in Spain, strawberry is a crop of high socioeconomic value that generates a large amount of direct and indirect jobs and incomes (López-Aranda *et al*. 2000). Especially considering that in 2021, Spain was the world’s largest exporter of strawberries, worth 853 million US dollars (Tridge, 2023). Particularly, the province of Huelva stands out with 87% of the total strawberry’s production in Spain (Instituto de Estadística y Cartografía de Andalucía, 2023). In Spain, strawberry plants are first grown in high altitude nurseries in provinces of Castilla y León, where optimal agroclimatic conditions to meet cold hour requirements prevail from March to September. Then, from October to June, runners are transplanted to production fields located in Huelva, province of Andalusia, where plants develop and fruit (Pastrana *et al*., 2017). This difference between the nursery and the production area not only provides ideal climatic conditions for the different growth stages, but also contributes as a phytosanitary measure helping to maintain the production area plagues free by breaking the life cycle of several pests and diseases. This contributes to the sustainability of the crop by reducing the use of necessary pesticides. In addition, one of the peculiarities about Huelva, a coastal area, is that there is an early production of the crops, allowing it to supply the market when there is not much stock of strawberries yet (Sáez Vera F. *et al*. 1996).

Given the importance of the strawberry cultivar on a socioeconomic level, this study aims to explore the metabolic processes which the strawberry plant undergoes during vernalization. To do so, a cutting-edge methodology, consisting in NMR-based metabolomic analysis in conjunction with computational tools for an automated holistic analysis of the metabolic profile and metabolite detection, is proposed. Furthermore, in this work, the potentialities of both NMR and analysis automation in the field of agribusiness are highlighted, contributing to enhancing precision, efficiency, and comprehensiveness in metabolomics studies through NMR.

## 2. Materials and methods

### 2.1. Experimental design and sample storage

Leaves from the plant mother and two daughters, to whom we will refer to as Node 1 (N1), Node 2 (N2), and Node 3 (N3), respectively, of 5 different strawberry plants (*Fragaria × ananassa ‘Viva Patricia’*) were collected at 6 different time points (every two weeks, from October 1 to December 16), frozen, and stored at -80 °C until metabolomic and proteomic analyses. The experimental design was validated through the calculation, in RStudio 1.4.1717 (RStudio Team, 2021), of the statistical power to detect large effects (Cohen’s d = 0.8) using assigned metabolomic data.

### 2.2. Sample preparation for Nuclear Magnetic Resonance

To obtain the soluble fraction, leaves were first frozen in liquid nitrogen and grinded using a mortar before alcoholic extraction. Afterwards, 50 mg of the powder leaf material was subjected to a 3-cycle process consisting of suspension of material in 2 mL of 80% ethanol in the first cycle, 50% ethanol in the second, and ultra-pure water in the last. Incubation for 15 min at 80 °C and subsequent cooling at 4 °C to diminish ethanol evaporation prior to centrifugation at 3000 g for 3 min was performed in each cycle. Supernatants obtained by centrifugation at the end of every cycle were kept, combined in one tube for each sample, and lyophilized for 24 h.

The freeze-dried material of each sample was dissolved in 540 µL of D_2_O 99.9%. Then, the aqueous extracts from the leaves were mixed with 60 µL of standardization buffer (0.2 M NaH_2_PO_4_, 5 mM of 3-(TrimethylSilyl) Propionic-2,2,3,3-d4 acid (TSP) and 30 mM NaN_3_, pH 7.0). A total 600 µL of the mixture was loaded into 5 mm NMR tubes (Bruker BioSpin srl) (Moing *et al*. 2004).

### 2.3. Nuclear Magnetic Resonance experiments

All the spectra shown in this work were acquired at the NMR Service of the CITIUS (https://citius.us.es/web/servicio.php?s=rmn), using a 500 MHz spectrometer. 1D-^1^H-Nuclear Overhauser Effect SpectroscopY (1D–^1^H–NOESY) with presaturation (d1) and spoil gradients (*noesygppr1d*.*2*) pulse sequence were used to acquire a ^1^H profile of samples while allowing water signal suppression (at ca. 4.8 ppm at 25 °C) to ease the identification of interesting signals at that same region (Parella *et al*. 2010). Mixing time, acquisition time, relaxation delay, and spectral width were 100 ms, 4 s, 4 s, and 20 ppm, respectively.

### 2.4. Nuclear Magnetic Resonance data preprocessing and quality control of spectra

Steps of chemical-shift referencing, phasing, baseline correction, and scaling of spectra according to the internal standard (TSP; ^1^H, 0.00 ppm) were performed in TopSpin 4.1.3. To ensure good spectral resolution, the TSP peak width at half height of each spectrum was checked to be ≤ 2.50 Hz for a line broadening of 1 Hz (Beckonert, 2007). Subsequently, spectra were locally aligned and peaks were detected in MATLAB R2021b (9.11.0.1769968) (MATLAB, 2021), using the package FOCUS (Alonso *et al*. 2014).

### 2.5. Metabolite identification and quantification of Nuclear Magnetic Resonance data

Spectra were assigned and quantified using two described methods. Firstly, in RStudio (RStudio Team, 2021), using the assignment and quantification algorithms, and the library of pure NMR spectra found in the R package Automatic Statistical Identification in Complex Spectra (ASICS). Also, using the Profiler module and library of the Chenomx NMR Suite V10.0 software (Chenomx Inc. Edmonton, Canada).

### 2.6. Metabolomic data analysis

Both untargeted and targeted analyses of NMR data were performed using RStudio tools (RStudio Team, 2021). Before analysis, due to the dynamic range of variables, data was normalized and centered. For the untargeted analysis, several descriptive methods of analysis were performed, including inspection of the global distribution before and after normalization, Pearson correlation calculation and representation in heatmaps, and Principal Component Analysis (PCA).

Regarding the targeted analysis, due to the independent, not normal distribution (*p* value of Shapiro-Wilk test < 10^-6^) and homoscedasticity of data (*p* value of Levene test > 0.01), Klustal Wallis test was performed to determine significant metabolites, and *p* values were adjusted for false positives using the Benjamini and Hochberg method. Only metabolites with adjusted *p* values lower than 0.1 were selected.

### 2.7. Protein extraction from strawberry leaves

For proteomic experiments, 10 frozen leaves were disrupted using a 6870 Freezer Mill (SPEX SamplePrep LLC) with 4 cycles of 1 min at 12 cycles per s and 1 min of cooling. After disruption, 250 mg of protein powder were resuspended in 1 mL of Extraction buffer (acetone 90%, TCA 10% + 8 mM DTT) and incubated at -20 °C overnight. After incubation, samples were centrifuged at 14000 g for 10 min at 4 °C and supernatants were discarded. Pellets were resuspended in 1 mL of Extraction buffer, disaggregated with 100 µL of glass beads, and further sonicated and centrifuged. Supernatants were discarded and pellets were washed several times with Extraction buffer until they became colorless. Finally, protein pellets were vacuum-dried and resuspended in Solubilization buffer (7 M urea, 2 M thiourea, 4% CHAPS). Protein samples were cleaned with 2D Clean-Up kit (Cytiva), and dissolved in rehydration buffer (4% CHAPS, 7 M urea, 2 M thiourea, 20 mM DTT, 0.5% IPG buffer 3-10 and 30 mM Tris-HCl pH 8.5). Total protein concentration was determined using RC DC™ Protein Assay (Bio-Rad).

### 2.8. 2D Difference Gel Electrophoresis

A total of 50 µg of protein from each sample were labeled with Cy3 or Cy5-Dye (Cytiva). Labeling was performed reciprocally so that each sample was separately labeled with Cy3 and Cy5 to account for any preferential protein labeling by the CyDyes. Also, 25 µg of each sample were labeled with Cy2-Dye and were pooled to use as an internal standard. Labeling was performed according to the manufacturer’s instructions. After labeling, samples were pooled. Isoelectric Focusing (IEF) was performed in 24 cm 3-10 NL Immobiline DryStrip (Cytiva) in an IPGphor unit (Cytiva) at 20 °C as follows: 10 h passive rehydration, 500 V for 1.5 h, linear gradient from 500 to 1000 V for 7 h, linear gradient from 1000 to 8000 V for 3 h and 8000 V for 5 h 36 min. During IEF, 62000 Vhr were reached. After IEF, strips were incubated for 15 min at room temperature in equilibration buffer (50 mM Tris-HCl pH 8.5, 30% v/v glycerol, 6 M urea, 2% w/v SDS) with 10 mg/mL DTT, followed by another 15-min incubation in equilibration buffer with 25 mg/mL iodoacetamide and loaded onto 12% acrylamide gels for the second dimension. The second dimension run was performed in an EttanDalt Six Electrophoresis Unit (GE Healthcare).

After electrophoresis, gels were imaged using Typhoon-9400 scanner (GE Healthcare) with a resolution of 100 μm using appropriate emission and excitation wavelengths, photomultiplier sensitivity, and filters for each of the Cy2, Cy3, and Cy5 dyes.

### 2.9. Protein profile quantification and differential analysis

Relative protein spot quantification across experimental conditions was performed using DeCyder v7.0 software and multivariate statistical module EDA v7.0 (Extended Data Analysis, GE Healthcare), as follows. First, the Batch Processor module detected spots from the three images of a gel (two experimental samples and one internal standard) and generated differential in-gel analysis images with information about spot abundance in each image with the value expressed as a ratio. After spot detection, the biological variation analysis module used those differential in-gel analysis images to match protein spots across all gels, using the internal standard for gel-to-gel matching. Statistical analysis was then carried out to determine protein expression changes. Spot presents in all gels and with abundance changes with a *p* value lower than 0.01 calculated from Student’s t-test with the multiple testing assessed using the false discovery rate were considered significant. Protein abundance changes were considered as relevant if they were ≤-5 or ≥5 with an ANOVA value lower than 10^-4^ where Component 1 was the node of the plant leaves and the Component 2 was the time of picking leaves. Multivariate analysis was performed by PCA using the algorithm included in the EDA module of the DeCyder v7.0 software based on the spots matched across all gels.

### 2.10. Protein identification

Protein identification was performed by in-gel trypsin digestion followed by MS at the Proteomics Service of the University Pablo de Olavide (Seville, Spain). Spots were excised from the gels manually and transferred to pierced V-bottom 96-well polypropylene microplates (BrukerDaltonik, Bremen, Germany) loaded with ultrapure water. Samples digestion and database searching Matrix-Assisted Laser Desorption/Ionization (MALDI)-MS and MS/MS were performed according to Shevchenko *et al*. 2006, with minor variations (Perkins *et al*. 1999; Schuerenberg *et al*. 2000; Suckau *et al*. 2003; Pérez-Pérez *et al*. 2009).

## 3. Results

### 3.1. Metabolic profile analysis of strawberry leaves reveals main sources of variability between samples

To understand the evolution of the strawberry plant metabolism throughout vernalization, leaves from 5 different strawberry plants at 6 different collection times (every two weeks from October 1 to December 16) were analyzed. Besides, to include possible variability between plants due to age, leaves were taken and analyzed separately from the plant mother and two daughters, to whom we will refer to as Node 1 (N1), Node 2 (N2), and Node 3 (N3), respectively.

At first sight, slight differences in terms of peak intensity between the metabolic ^1^H-NMR profiles of the three nodes were observed (**Figure 1A**), suggesting that the age of the plant was not the main source of variability among samples. These similarities between the metabolic profiles of the three nodes were further verified by PCA. As can be seen in **Figure 1B**, the two main components (Principal Component 1, PC1; and Principal Component 2, PC2) which explain, respectively, 28.5% and 17% of the total variance, and together, comprised almost half the total variability of the dataset (45.5%), were not able to separate samples from N1, N2, and N3 of all the collected time points in individual groups. Hence, considering that there were no relevant changes in the metabolic profile of samples due to age differences, samples of different nodes were considered replicates in subsequent analyses. In other words, samples from three nodes at a specific time point were considered three biological replicates.

**Figure 1.**
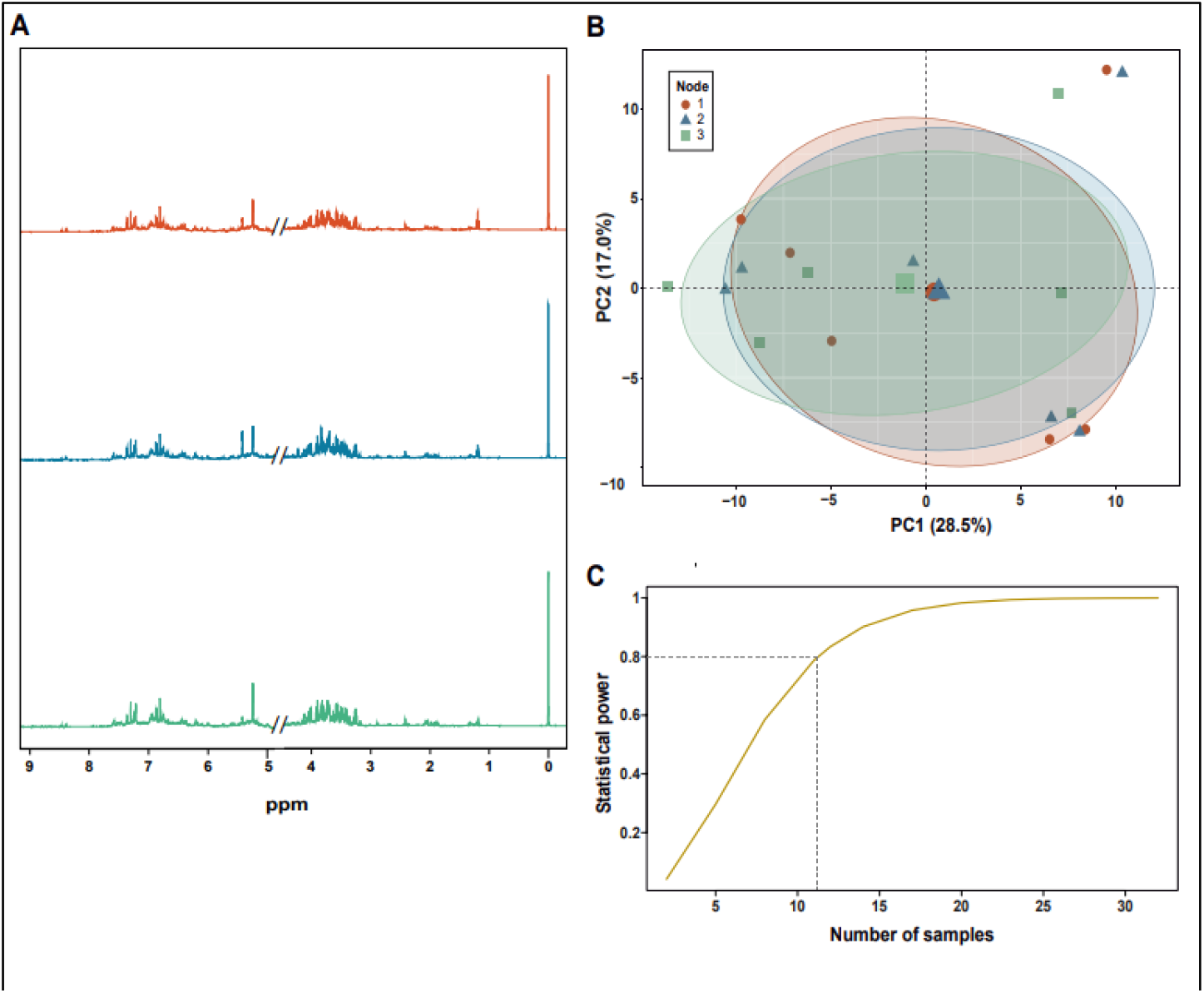
Study of strawberry leaves from different plants according to age. **A)** Comparison of ^1^H-NOESY; noesygppr1d spectra aliphatic and aromatic regions (1-3 and 5-9 ppm, respectively) of strawberry leaf samples of three different nodes (N1: Node 1; N2: Node 2; N3: Node 3, colored in red, blue, and green, respectively) collected on October 1. All spectra are referenced according to the chemical shift reference frequency provided by the TSP signal (^1^H, 0.00 ppm). The aromatic region was scaled x4 with respect to the rest of the spectra just for figure representation, to distinguish low-intensity signals more clearly. **B)** Principal Component Analysis (PCA) of normalized NMR data from strawberry leaves grouped by node. Each point corresponds to one NMR spectrum acquired for one sample and is colored according to the type of node. For each class of sample, the 95% confidence ellipse with its corresponding centroid (biggest point in the center of each ellipse) is shown. Sample labels and color key are the same as in (A). **C)** Graph representing the statistical power of the experimental design *versus* the number of samples of the experiment. Using gray dotted lines, the statistical power threshold and the corresponding number of samples required to achieve it are indicated.

Then, to evaluate the suitability of the experimental design proposed, the statistical power was calculated. As it is shown in **Figure 1C**, 12 samples were enough to detect large effects with sufficient statistical power (at least 0.8), and thus, the experimental design proposed (18 samples) was adequate for the purpose of the study.

Thereafter, when inspecting the NMR-based metabolic profile of samples collected at different time points, differences were easily noticeable (**Figure 2A**). For additional confirmation, using PCA, it was possible to corroborate that changes among samples came predominantly from collection time points, which are set apart in different groups in **Figure 2B**, independently of the node. The first group of samples with similar metabolic profiles included samples from October 1 to October 30, as their confidence ellipses overlapped. Then, a second group with samples from November 14 appeared. Finally, two closely located but independent groups existed for samples from December 3 and December 16. The distribution of these groups can be explained by analyzing the plot axis in more detail, since along the PC1 or x-axis, samples were segregated in early collection time points (from October 1 to October 30; negative values of the PC1) and late collection time points (from November 14 to December 16). Concerning the PC2, samples from early collection time points appeared to be scarcely influenced by this variable, since they were located at the y-axis zero. Instead, this PC separated samples collected on November 14 (positive values of the PC2) from those collected after that day, namely, December 1 and 16 (negative values of the PC2).

**Figure 2.**
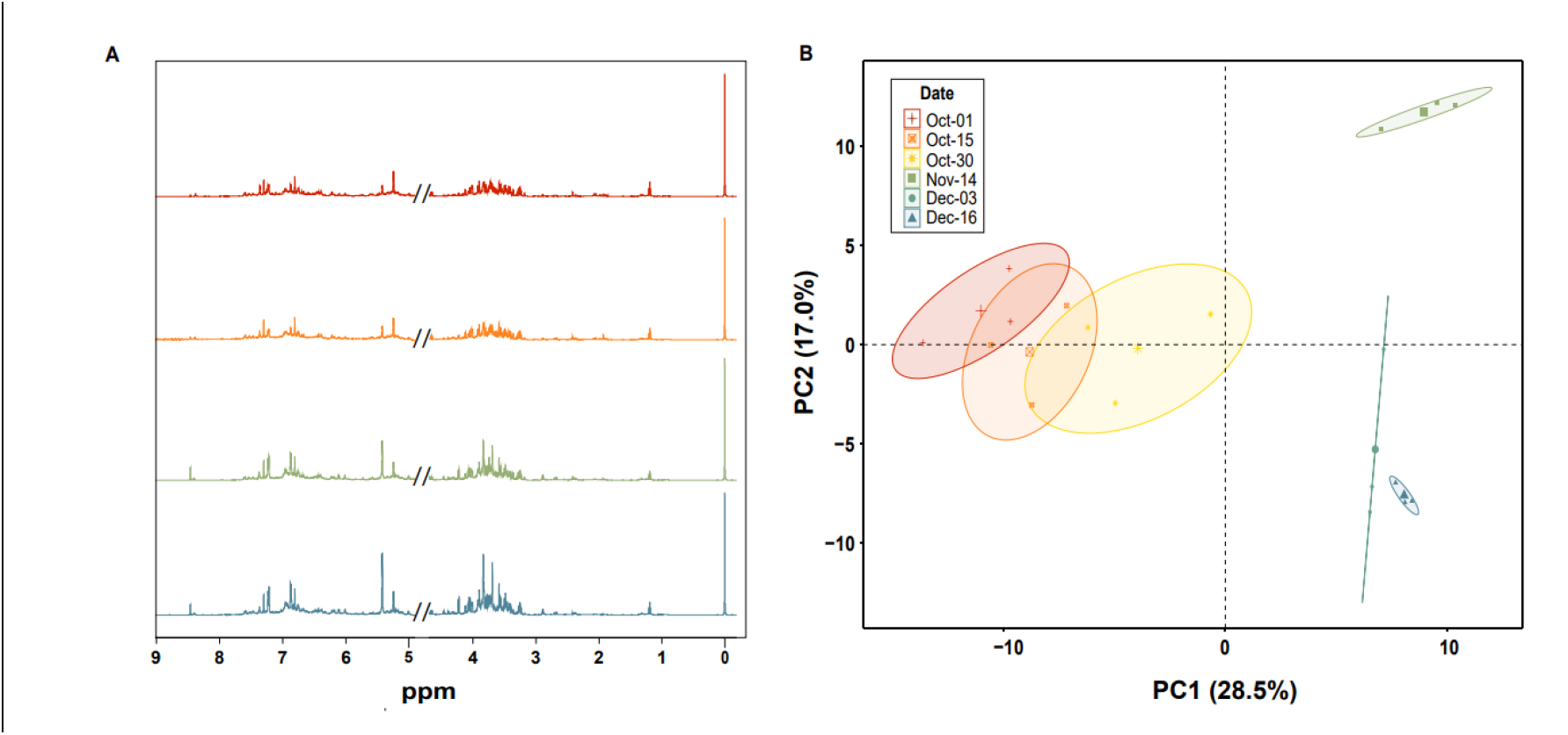
Influence of collection time point in strawberry leave metabolic profile. **A)** Comparison of ^1^H-NOESY; noesygppr1d spectra aliphatic and aromatic regions (1-3 and 5-9 ppm, respectively) of strawberry leaf samples collected at 4 time points (October 1, October 15, November 14, and December 16). All spectra are referenced according to the chemical shift reference frequency provided by the TSP signal (^1^H, 0.00 ppm). The aromatic region was scaled x4 with respect to the rest of the spectra just for figure representation, to distinguish low-intensity signals more clearly. **B)** Principal Component Analysis (PCA) of normalized NMR data from strawberry leaves grouped by time of collection. Each point corresponds to one NMR spectrum acquired for one sample and is colored using a gradient from warm to cool colors according to the time of collection (from October 1 to December 16). For each class of sample, the 95% confidence ellipse with its corresponding centroid is shown. PC1 or x-axis segregates samples in early collection time points (from October 1 to October 30; negative values of the PC1) and late collection time points (from November 14 to December 16). PC2 or y-axis scarcely influences samples from early collection time points, and segregates samples collected on November 14 (positive values of the PC2) from later times of collection, December 1 and 16 (negative values of the PC2).

It is important to note that the previous analyses were performed on normalized data. Due to the fact that NMR is inherently proportional to sample concentration and that metabolites can be found in a wide dynamic range of concentrations in comparison to other types of biomolecules, data normalization is useful to weight signals and variables for importance and relevance in the study context, rather than based only on abundance or intensity (Vignoli *et al*. 2019). In this sense, global distribution analysis before normalization (**Figure S1A**) showed that NMR spectra of strawberry leaves were mainly made up by abundant low-intensity signals, and few high-intensity signals. Also, when performing sample correlation analyses before normalization (**Figure S1B**), values of correlation coefficients among samples were moderate, and showed the presence of two main groups: one with samples collected on early time points (from October 1 to October 30), and another with samples collected on late time points (from November 14 to December 16).

After normalization, the global distribution of samples was modified. Firstly, the intensities range was decreased, and therewith, the sample quartiles became more balanced (**Figure S1C**). When Pearson correlation was calculated over normalized data, sample correlations were generally increased, and different, more defined sample groups could be identified. Specifically, three groups could be distinguished: one with samples collected on October 1, 15, and 30; a second one with samples collected on November 14; and, the last one, with samples collected on December 3 and 16 (**Figure S1D**). Hence, patterns found on heatmaps before and after normalization agreed with the differences per collection time point observed previously in NMR spectra and PCA (**Figure 2**). This highlights the importance of data treatment in reaching relevant conclusions in the context of massive data analysis.

### 3.2. Metabolite annotation allows understanding of changes in metabolic profile

Given that the first and main source of changes in the groups of metabolic profiles (PC1) appeared in between the group of samples collected from vernalization in October (1, 15, and 30) and November 14, and that assignment is computational and time-consuming, we focused on studying this first step in the evolution of the metabolic profile. Purposely, spectra from October 15 and November 14 were used for automatic assignment, reaching up to 103 putatively annotated compounds, of which 62 changed significantly (adjusted *p* value < 0.1). After that, the automated annotation was checked by comparing the chemical shifts of peaks with reference spectra from the Human Metabolome Database (HMDB).

The expression patterns of three annotated metabolites from October 15 to November 14 are shown in **Figure 3**; two of them, Glucose 6-phosphate (G6P) and gamma-aminobutyric acid (GABA) levels decrease, whereas the levels of sucrose and fumarate, increase from October 15 to November 14. G6P is a molecule commonly found in living beings, mainly related to processes from the energetic metabolism, such as glycolysis, or the pentose phosphate pathway. This molecule has many possible fates in cell metabolism, and can act as substrate for storage in the form of polysaccharides -starch in the case of plants-. Regarding GABA, it is a conserved molecule among species with structural, energetic and signaling functions. Concerning sucrose, this sugar is involved in energetic metabolism, signaling and stress adaptation. Finally, in relation to fumarate, it is a carboxylic acid from the tricarboxylic acid cycle (TCA), being also involved in the urea cycle.

**Figure 3.**
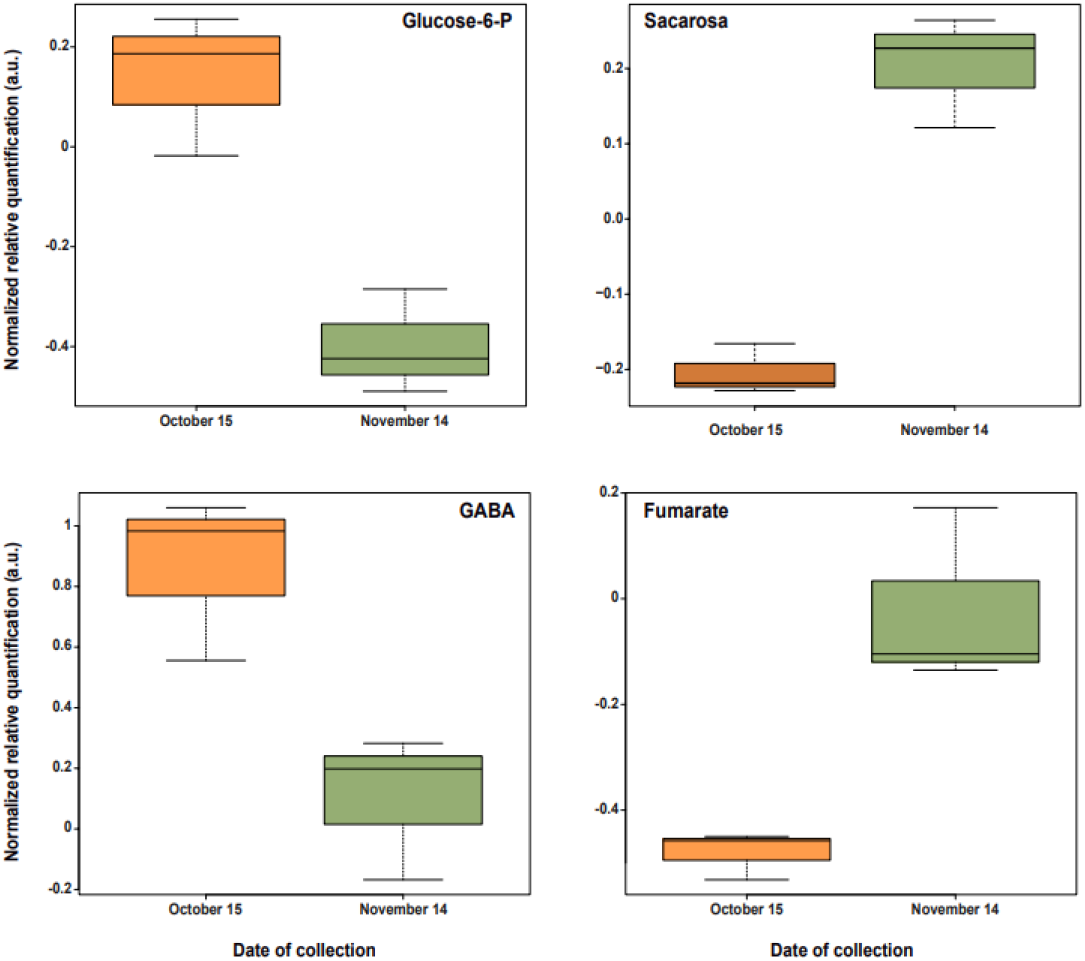
Boxplots of metabolomic annotation from NMR data of samples collected on October 15 and November 14. From *left* to *right*, boxplots for glucose 6-phosphate, GABA, and fumarate. Horizontal lines of each box show the *upper* quartile, median, and *lower* quartile. Also, maximum and minimum values are represented as horizontal lines in the extremes of the whiskers (vertical dotted lines).

### 3.3. Protein profile analysis of strawberry leaves supports NMR-based metabolic profile analysis

To understand the origin of the distinguished metabolic profiles, proteomic analysis was carried out. Considering that, after performing metabolomic analysis, differences between nodes were minimum compared to the time of collection, only two nodes were used for the analysis, the oldest and the youngest, N1 and N3, respectively. Also, as changes in proteomic profile are expected to be slower than those regarding metabolites, only proteomic analysis of the first and last times of collection (September and December, respectively) were used for comparison by 2D Difference Gel Electrophoresis (2D-DIGE).

Three replicas of the four types of samples were run in eight independent gels, each containing two experimental samples labeled with Cy3 and Cy5 and an internal standard comprised of pooled samples from all strains labeled with Cy2. The presence of the internal standard allowed the normalization of individual protein abundance between samples from different gels. Using DeCyder v7.0 software, we could detect 767 protein spots present in 100% of gel images. From those spots, 525 had an individual protein abundance highly reproducible between replicate samples from the time of recollection condition (two-way ANOVA condition 2 < 0.001) and 12 were selected for further identification by MS with changes in protein abundance of more than fivefold between September and December and could be cut from the acrylamide gel (**Figure S2A-B**). Using PCA, we showed that the two first components could explain the 97% of differences between samples, where PC1 accumulated 90.6% of the variance, and separated samples by collection time point, whereas PC2 only comprised 6.4% (**Figure S2C**). These results are in line with previous metabolomic analysis (**Figure 1**), and thus confirm the applicability of NMR untargeted analysis in plant and agricultural contexts.

### 3.4. Differences in protein expression are found between early and late collection time points

From the 12 spots analyzed using MALDI–MS or MALDI MS/MS, identification was successfully obtained for 10 spots, corresponding to 8 different proteins (**Table 1**). Spots #1, #2, #3, #6, #7, #8, #9, #10, and #12 were found over-expressed in December, and, with the exception of spot #10, they all correspond to proteins involved in light-independent stages of photosynthesis, also known as the Calvin cycle. Particularly, 3 spots were proteins of the Ribulose-1,5-Bisphosphate Carboxylase/Oxygenase (RuBisCO, EC 4.1.1.39) -key enzyme of the Calvin Cycle, responsible for carbon fixation-; spots #2 and #3 were identified as large chains of that same enzyme, and spot #6 corresponds to small chain 1. Spot #8 is RuBisCO large subunit-binding protein subunit beta (Table 1) (Ellis & Van Der Vies, 1988).

**Table 1.**
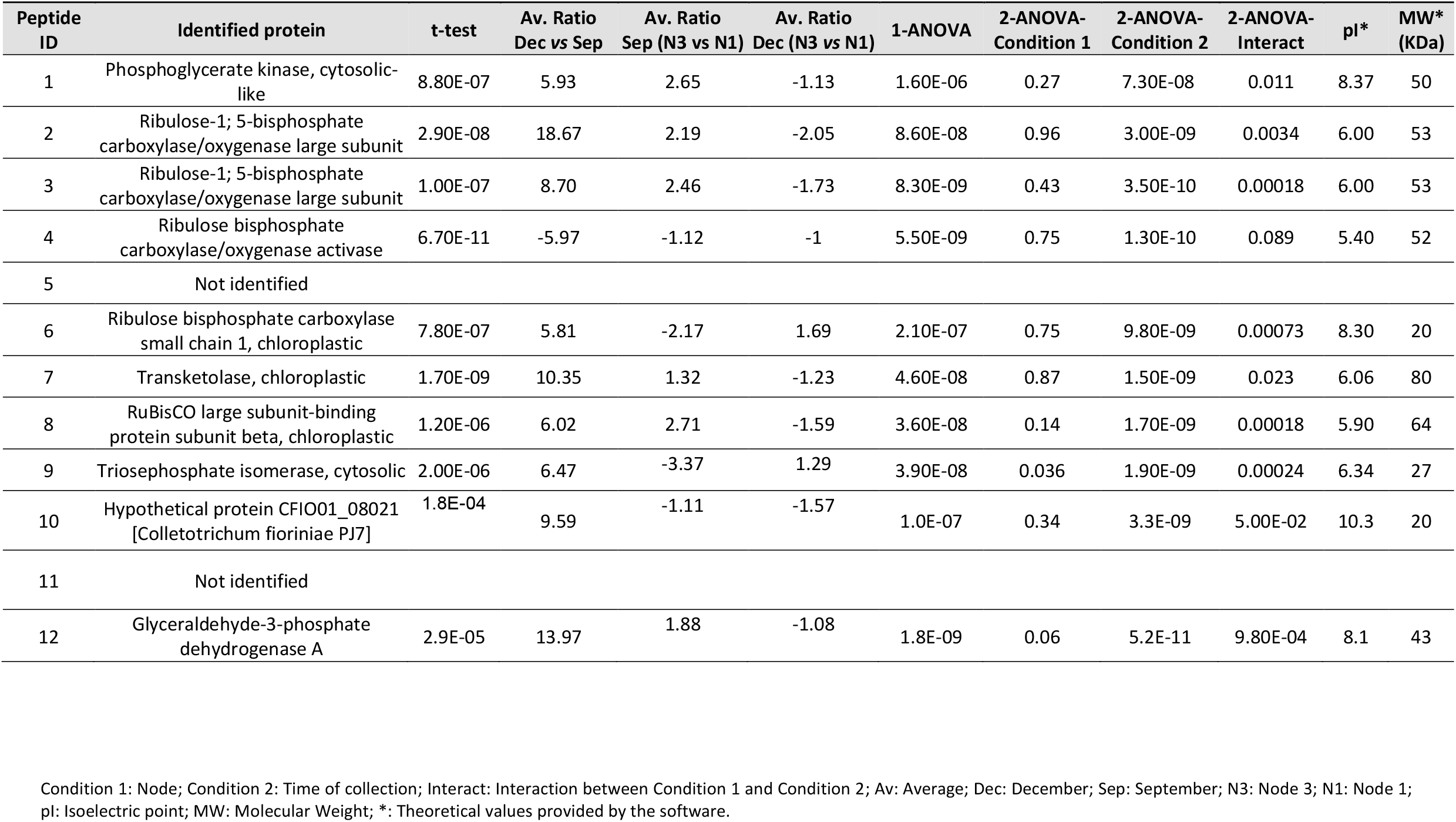
Protein identification by MALDI-MS and MS/MS.

| Peptide ID | Identified protein | t-test | Av. Ratio Dec vs Sep | Av. Ratio Sep (N3 vs N1) | Av. Ratio Dec (N3 vs N1) | 1-ANOVA | 2-ANOVA-Condition 1 | 2-ANOVA-Condition 2 | 2-ANOVA-Interact | pI* | MW* (kDa) |
| --- | --- | --- | --- | --- | --- | --- | --- | --- | --- | --- | --- |
| 1 | Phosphoglycerate kinase, cytosolic-like | 8.80E-07 | 5.93 | 2.65 | -1.13 | 1.60E-06 | 0.27 | 7.30E-08 | 0.011 | 8.37 | 50 |
| 2 | Ribulose-1; 5-bisphosphate carboxylase/oxygenase large subunit | 2.90E-08 | 18.67 | 2.19 | -2.05 | 8.60E-08 | 0.96 | 3.00E-09 | 0.0034 | 6.00 | 53 |
| 3 | Ribulose-1; 5-bisphosphate carboxylase/oxygenase large subunit | 1.00E-07 | 8.70 | 2.46 | -1.73 | 8.30E-09 | 0.43 | 3.50E-10 | 0.00018 | 6.00 | 53 |
| 4 | Ribulose bisphosphate carboxylase/oxygenase activase | 6.70E-11 | -5.97 | -1.12 | -1 | 5.50E-09 | 0.75 | 1.30E-10 | 0.089 | 5.40 | 52 |
| 5 | Not identified |  |  |  |  |  |  |  |  |  |  |
| 6 | Ribulose bisphosphate carboxylase small chain 1, chloroplastic | 7.80E-07 | 5.81 | -2.17 | 1.69 | 2.10E-07 | 0.75 | 9.80E-09 | 0.00073 | 8.30 | 20 |
| 7 | Transketolase, chloroplastic | 1.70E-09 | 10.35 | 1.32 | -1.23 | 4.60E-08 | 0.87 | 1.50E-09 | 0.023 | 6.06 | 80 |
| 8 | RuBisCO large subunit-binding protein subunit beta, chloroplastic | 1.20E-06 | 6.02 | 2.71 | -1.59 | 3.60E-08 | 0.14 | 1.70E-09 | 0.00018 | 5.90 | 64 |
| 9 | Triosephosphate isomerase, cytosolic | 2.00E-06 | 6.47 | -3.37 | 1.29 | 3.90E-08 | 0.036 | 1.90E-09 | 0.00024 | 6.34 | 27 |
| 10 | Hypothetical protein CFIO01_08021 [Colletotrichum fioriniae PJ7] | 1.8E-04 | 9.59 | -1.11 | -1.57 | 1.0E-07 | 0.34 | 3.3E-09 | 5.00E-02 | 10.3 | 20 |
| 11 | Not identified |  |  |  |  |  |  |  |  |  |  |
| 12 | Glyceraldehyde-3-phosphate dehydrogenase A | 2.9E-05 | 13.97 | 1.88 | -1.08 | 1.8E-09 | 0.06 | 5.2E-11 | 9.80E-04 | 8.1 | 43 |

Condition 1: Node; Condition 2: Time of collection; Interact: Interaction between Condition 1 and Condition 2; Av: Average; Dec: December; Sep: September; N3: Node 3; N1: Node 1; pI: Isoelectric point; MW: Molecular Weight; \*: Theoretical values provided by the software.

Likewise, spot #1 was identified as the enzyme phosphoglycerate kinase (EC 2.7.2.3), which intervenes right after the carboxylation step of RuBisCO as part of the regeneration process of ribulose-1,5-bisphosphate (RuBP). Subsequent enzymes of the Calvin cycle, similarly involved in RuBP regeneration, were identified in the remaining mentioned spots, being spot #12 the glyceraldehyde-3-phosphate dehydrogenase (EC 1.2.1.12), spot #9 the triphosphate isomerase (EC 5.3.1.1), and spot #7 a transketolase (EC 2.2.1.1).

Regarding spot #10, it was also over-expressed in December, and it was identified as a hypothetical protein CFIO01_08021 from the plant pathogen fungi, *Colletotrichum fioriniae* PJ7. This fungus has been proven to cause anthracnose in strawberry fruits, as well as root necrosis. However, besides decreasing strawberry quality and production, this pathogen is not harmful for humans. Spots 5 and 11 could not be identified.

Only spot number #4, from all the analyzed by MS and MS/MS, was found over-expressed in September, and matched with the Ribulose-1,5-bisphosphate Carboxylase/oxygenase Activase (RCA, EC 4.1.1.36), an enzyme whose function is the activation of RuBisCO.

## 4. Discussion

Vernalization is a physiological process of plants of great interest, not only from a research point of view, but also from industrial and economic ones, due to its close relation with flowering and fruit production. This explains the wide variety of studies dedicated to comprehending the mechanisms driving it, and that go from genetic, physiological and structural approaches of classical biochemistry (Velanis *et al*. 2017; Amasino, 2004), to, in more recent years, study of the whole composition of biomolecules, namely, omics technologies such as transcriptomics and metabolomics (Villacorta-Martin *et al*. 2015; Lv *et al*. 2022; Chen *et al*. 2021). However, the vast majority of these metabolomic studies are performed using MS rather than NMR, even though this technique has previously been proven useful for metabolite profile analysis in diverse application fields, including biomedicine and food industry (Wu *et al*. 2022; Chowdhury *et al*. 2023).

The main reasons behind this preference may be related with the complexity of ^1^H-NMR spectra; as they are composed by thousands of overlapping peaks, visual and manual analyses are time consuming, biased, and inadequate. This was shown previously in **Figure 1A**, where even though there may be small differences between strawberry nodes when looking at the NMR spectra, subsequent automated analysis showed, in **Figure 1B**, that these differences were not as relevant as they may seem. In this sense, data preprocessing, and especially normalization have also shown to be essential in reaching accurate conclusions in an objective and automated manner, and support the idea that visual inspection of raw NMR spectra is insufficient.

From both the untargeted and targeted analyses of metabolomic and proteomic data, it becomes clear that there is an evolution of strawberry plants throughout vernalization, which is the main source of variability between samples. In particular, the metabolic profile of strawberry leaves shows that, in early times of collection, such evolution is slow and light, resulting in no statistically relevant differences between samples from October 1 to October 30. However, during the following 15 days, to November 14, a drastic change is appreciated. We propose that this greater effect may be related to vernalization. In vernalization, plants develop the ability to flower and produce fruits after going through a dormant period as a consequence to cold exposure. Thus, it would be expectable that, at the beginning of vernalization, the metabolic profile of plants remains relatively stable, until sufficient cold hours are accumulated. Then, the plant would activate its metabolism to get ready for flowering and fruit production, being both high energy demanding processes.

To understand the implications of the metabolomic and proteomic profile changes in samples, targeted analyses of both metabolomic and proteomic data were carried out. Protein identification and differential analysis suggest that in early collection time points, the strawberry plants may be still in dormancy -given that metabolic profile evolution is slow as shown in **Figure 2**- but getting ready for posterior processes that may require energy, hence the activation of the RCA enzyme (**Table 1**). Then, in late collection time points, the enzymes of the Calvin cycle are overexpressed, including the RuBisCO, which is consistent with previous activation of RCA. At this point, cells ought to meet the energy requirements to continue their development.

Looking at the annotated metabolites, it is possible to get an idea of the functions and metabolic pathways that are favored throughout the vernalization process. Notably, the assigned compounds found in **Figure 3** imply that vernalization effects on plant metabolism occur at different levels. First, in energetic metabolism and storage, where it has been described that a decrease in G6P (**Figure 4**) -a substrate for energy storage as starch-may be related to an increment in sugar levels. Those sugars can furtherly be used by the plant for energetic purposes, including fruit production and ripening (Preiser *et al*. 2018). This agrees with the overexpression of Calvin cycle proteins for carbon fixation, as well as with the increase observed for sucrose, which is the most common sugar found in the phloem sap of many plants, since it is cheap to produce and easily available (Thomson & Thorpe, 1987) (**Figure 4**)

**Figure 4.**
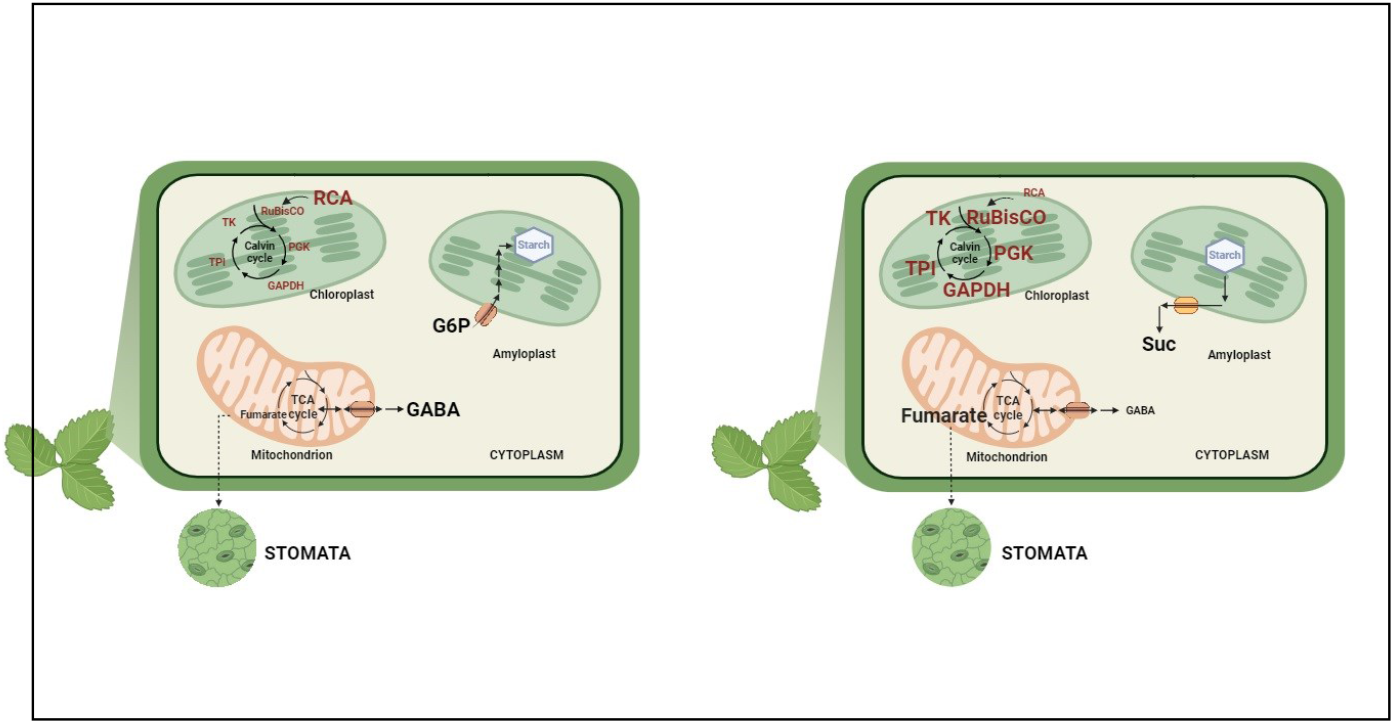
Model behavior for plant metabolism in early and late times of vernalization. Two models for plant cells of strawberry leaves are proposed depending on the stage of vernalization. In early collection time points, a Dormant metabolism cell is found with starch and GABA synthesis activated, as well as RCA enzyme expression, preparing the cell for future high demanding processes. In late collection time points, an Active metabolism cell starts, with TCA and Calvin cycle functioning. Moreover, fumarate produced in TCA, may leave the current cell to attend cell guards and regulate stomata movement. Metabolites are represented using plain colors; for proteins, a striped pattern was used. Deactivated processes are shown in gray, whereas activated processes are colored. G6P: glucose 6-phosphate; GABA: gamma-aminobutyrate; GADPH: glyceraldehyde-3-phosphate dehydrogenase; RCA: ribulose bisphosphate carboxylase/oxygenase activase; RuBisCO: Ribulose Bisphosphate Carboxylase/Oxygenase; TCA: tricarboxylic Acid; TPI: triphosphate isomerase; TK: transketolase; PGK: phosphoglycerate kinase.

Moreover, the GABA molecule has been found to be related with survival in stress conditions through plant signaling and homeostatic functions. Previous studies in tomato (Li *et al* 2021) describe that increasing GABA levels inhibit plant growth, delay and reduce flowering, and lessen fruit yield. In the vernalization context, a decrease in GABA levels as we found for strawberry plants, might be an indication of successful flowering and fruit bearing. In addition, it has been demonstrated that GABA plays a role in soothing the effects of cold stress, such as cytoplasmatic acidosis and the generation of reactive oxygen species (ROS). This is achieved through proton consuming reactions involved in its synthesis, which regulates cytosolic pH; and its ROS-scavenging capacities. Finally, as cold stress lessens, GABA is turned into a Krebs cycle intermediate, enhancing plant metabolism (**Figure 4**) (Fusco *et al*. 2023).

In plan metabolism, fumarate has been observed to serve as a secondary storage molecule when starch availability is limited, such as during reproductive growth. Its contribution lies in its capacity to act as a carbon sink, supplying energy needs during growth through carbon mobilization (Olas *et al*. 2020). Besides being an indicator of energetic metabolism as its function as part of the TCA cycle, has a new function as a structure and movement regulator of stomata (**Figure 4**) (Medeiros *et al*. 2015). Several studies propose that the function of fumarate in stomata is similar to that of malate (Araújo *et al*. 2011) and have proven to correlate fumarate levels to stomatal kinetics (Lemonnier *et al*. 2023), which supports the impact of this molecule in stomata movements. In this sense, metabolomic studies are being used to study stomatal movements (Medeiros *et al*. 2019), although they mainly use MS. In **Figure 4**, a model for leaves cell behavior in early and late times of collection is proposed.

Taken together, NMR-based metabolomic studies, in conjunction with automation tools, emerge as an interesting option to study the evolution and progress of crops, as has been done in this study using strawberry, demonstrating that it is able to capture subtle changes in plant development. Future work may be useful to address some remaining questions. For instance, including a higher number of samples could improve the statistical power, to detect not only large effects, but also to reach subtle changes that may be of relevance. Also, it would be interesting to gather metadata regarding viability and productivity, to be able to correlate changes in metabolic profile to parameters of use for cultivars, thus making out of NMR-based metabolomic analysis a crop monitoring tool. In all cases, this work surely opens new avenues for the use of NMR in an agricultural context.

## Acknowledgements

This work was supported by the Spanish Government (PID2021-126663NB-I00 funded by MCIN/AEI/ 10.13039/501100011033 and by ERDF A way of making Europe to I.D.-M.), European Regional Development Fund (FEDER).

## Author contributions

All authors have made a substantial, direct and intellectual contribution to the work, and approved it for publication.

## References

Abreu A. C. and Fernández I., (2020) NMR Metabolomics Applied on the Discrimination of Variables Influencing Tomato (Solanum lycopersicum). Molecules 25, 3738.

Alonso A., Marsal S., and Julia A., (2015) Analytical Methods in Untargeted Metabolomics: State of the Art in 2015. Front. Bioeng. Biotechnol. 3.

Amasino R. (2004). Vernalization, Competence, and the Epigenetic Memory of Winter. Plant Cell 16, 2553–2559.

Araújo W. L., Nunes-Nesi A. and Fernie A. R. (2011). Fumarate: Multiple functions of a simple metabolite. Phytochem. 72, 838–843.

Bahukhandi A., Attri D.C., Mehra T. and Bhatt I.D., (2022) Himalayan Fruits and Berries: Bioactive Compounds, Uses and Nutraceutical Potential, Elsevier, Amsterdam, pp. 183–196.

Instituto de Estadística y Cartografía de Andalucía (2023, November) Boletín de Coyuntura Provincial [Regional Juncture Newsletter] https://www.juntadeandalucia.es/institutodeestadisticaycartografia/boletincoyuntura/boletin.htm?p=21

Chen J., Xue M., Liu H., Fernie A. R. and Chen W. (2021) Exploring the genic resources underlying metabolites through mGWAS and mQTL in wheat: From large-scale gene identification and pathway elucidation to crop improvement. Plant Commun. 2(4).

Chowdhury C. R., Kavitake D., Jaiswal K. K., Jaiswal K. S., Reddy G. B. et al. (2023). NMR-based metabolomics as a significant tool for human nutritional research and health applications. Food Biosci. 53, 102538.

Costa R. C. da., Calvete E. O., Mendonça H. F. C. and DeCosta L. A., (2014) Phenology and leaf accumulation in vernalized and non-vernalized strawberry seedlings in neutral days. Acta Sci. Agron. 36, 57–62.

Ellis R. J., Van Der Vies S. M., (1988) The Rubisco subunit binding protein. Photosynth Res. 16(1-2):101–115

Emwas A. H., Roy R., McKay R. T., Tenori L., Saccenti E. et al., (2019) NMR Spectroscopy for Metabolomics Research. Metabolites 9(7), 123.

Fusco G. M., Carillo P., Nicastro R., Pagliaro L., De Pascale S. et al., (2023) Metabolic profiling in tuberous roots of Ranunculus asiaticus L. as influenced by vernalization procedure. Plants 12, 3255.

García-Villaescusa A., Morales-Tatay J.M., Monleón-Salvadó D., González-Darder J.M., Bellot-Arcis C. et al., (2018) Using NMR in saliva to identify possible biomarkers of glioblastoma and chronic periodontitis. PLoS ONE 13(2).

Gavaghan C. L., Holmes E., Lenz E., Wilson I. D. and Nicholso J. K., (2000) An NMR-based metabonomic approach to investigate the biochemical consequences of genetic strain differences: application to the C57BL10J and Alpk:ApfCD mouse. FEBS Lett. 484, 1873–3468.

Wishart D. S., Guo A. C., Oler E. et al. (2022). HMDB 5.0: the Human Metabolome Database for 2022. Nucleic Acids Res. 50:D622.

Johnson C. H. and Gonzalez F. J., (2018) Challenges and opportunities of metabolomics. J. Cell. Physiol. 227, 2975.

Lemonnier P. and Lawson T. (2023). Seminars in Cell and Developmental Biology (Calvin cycle and guard cell metabolism impact stomatal function).

Li L., Dou N., Zhang H. and Wu C. (2021). The versatile GABA in plants. Plant Signal. Behav. 16(3).

López-Aranda J. M., Bartual R., Marsal J. I., Medina J. J., Lopez-Montero R. et al. (2000) Panorama varietal de la fresa en España. Los programas de obtención de nuevas variedades. [Strawberry varietal outlook in Spain. Breeding new varieties programs.] Agrícola vergel 218, 82–92.

Lv L., Dong C., Liu Y., Zhao A., Zhang Y. et al. (2022). Transcription-associated metabolomic profiling reveals the critical role of frost tolerance in wheat. BMC Plant Biol. 22, 333.

MATLAB, 2021. version 9.11.0.1769968 (R2021b), Natick, Massachusetts: The MathWorks Inc.

Medeiros D. B., Daloso D. M., Fernie A. R., Nikoloski Z., and Araújo W. L. (2015). Utilizing systems biology to unravel stomatal function and the hierarchies underpinning its control. Plant Cell Environ. 38, 1457.

Medeiros D. B., Luz L. M. da, Oliveira H. O. da, Araújo W. L., Daliso D. M. et al. (2019). Metabolomics for understanding stomatal movements. Theor. Exp. Plant Physiol. 31, 91.

Moco S., (2022) Studying Metabolism by NMR-Based Metabolomics. Front. Mol. Biosci. 9.

Moing A., Maucourt M., Renaud C., Gaudillère M., Brouquisse R. et al. (2004). Quantitative metabolic profiling by 1-dimensional 1H-NMR analyses: application to plant genetics and functional genomics. Funct. Plant Biol. 31(9), 889–902.

Morales A., Carmen Gloriay Vargas S., Sigrid, Eds. (2017) Manual de manejo agronómico de la Frutilla [Manual for strawberry agronomic management], Boletín INIA Nº 17 - Instituto de Investigaciones Agropecuarias, Villa Alegre.

Olas J. J., Apelt F., Watanabe M., Hoefgen R. and Wahl V., (2020) Developmental stage-specific metabolite signatures in Arabidopsis thaliana under optimal and mild nitrogen limitation. Plant sci. 303, 110746

Oviedo V. R. S., Enciso-Garay C. R. and García Figueredo E. I., (2020) Vernalizing pre-transplants improved the agronomic characteristics of strawberry genotypes under tropical conditions. Rev. Caatinga. 33, 653–659.

Parella, T. (2010). Pulse Program Catalogue: I. 1D & 2D NMR EXPERIMENTS. Accessed 23 May 2021, from http://www.biocore.pku.edu.cn/public/Uploads/file/20170901/20170901085426_59631.pdf

Pastrana, A. M., Basallote-Ureba M. J., Aguado A., and Capote N. (2017). Potential Inoculum Sources and Incidence of Strawberry Soilborne Pathogens in Spain. Plant Dis. 101, 751–760.

Pérez-Pérez R., Ortega-Delgado F. J.,García-Santos E., López J. A., Camafeita E. et al. (2009). Differential Proteomics of Omental and Subcutaneous Adipose Tissue Reflects Their Unalike Biochemical and Metabolic Properties. J. Proteome Res. 8(4), 1682–1693

Perkins D. N., Pappin D. J. C., Creasy D. M. and Cottrell J. S. (1999). Probability-based protein identification by searching sequence databases using mass spectrometry data. Electrophoresis 20, 3551–3567.

Preiser A. L., Banerjee A., Fisher N., and Sharkey T. D. (2018). BioRxiv 442434 (Supply and consumption of glucose 6-phosphate in the chloroplast stroma)

RStudio Team (2021). RStudio: Integrated Development Environment for R. RStudio, PBC, Boston, MA. Retrieved from http://www.rstudio.com/

Sáez Vera F. and Ibáñez Pelayo L., (1996) Los viveros de fresa en castilla y león. Produccion de plantas jovenes de fresón para Huelva. Agricultura 768, 592–595.

Schuerenberg M., Luebbert C., Eickhoff H., Kalkum M., Lehrach H. et al. (2000). Prestructured MALDI-MS Sample Supports. Anal. Chem. 72(15), 3436–3442.

Shevchenko A., Tomas H., Havli J., Olsen J. V. and Mann M., (2006). In-gel digestion for mass spectrometric characterization of proteins and proteomes. Nat. Protoc. 1, 2856–2860.

Suckau D., Resemann A., Schuerenberg M., Hufnagel P., Franzen J. et al. (2003). A novel MALDI LIFT-TOF/TOF mass spectrometer for proteomics. Anal. Bioanal. Chem. 376, 952–965.

Thomson M., Thorpe T. A., Bong J. M., Durzan D. J., Eds. (1987) Cell and tissue culture in forestry, Martinus Nijhoff Publishers, Dordrecht, pp 89–112

Tridge. (2023, November) Fresh Strawberry. Tridge. https://www.tridge.com/es/intelligences/stawberry

Velanis C. N., and Goodrich J. (2017). Vernalization and Epigenetic Inheritance: A Game of Histones. Curr. Biol. 27, 1324.

Vignoli A., Ghini V., Meoni G., Licari C., Takis P. G. et al. (2019). High-Throughput Metabolomics by 1D NMR. Angew. Chem. Int. Ed. 58, 968.

Villacorta-Martin C., Núñez de Cáceres González F. F., de Haan J., Huijben K., Passarinho P. et al. (2015). Whole transcriptome profiling of the vernalization process in Lilium longiflorum (cultivar White Heaven) bulbs. BMC Genom. 16, 550.

Wu W., Zhang L., Zheng X., Huang Q., Farag M. A. et al. (2022). Emerging applications of metabolomics in food science and future trends. Food Chem.: X 16(30), 100500.

