## Supplemental Figures for "Applying an automated NMR-based metabolomic workflow to unveil strawberry molecular mechanisms in vernalization"

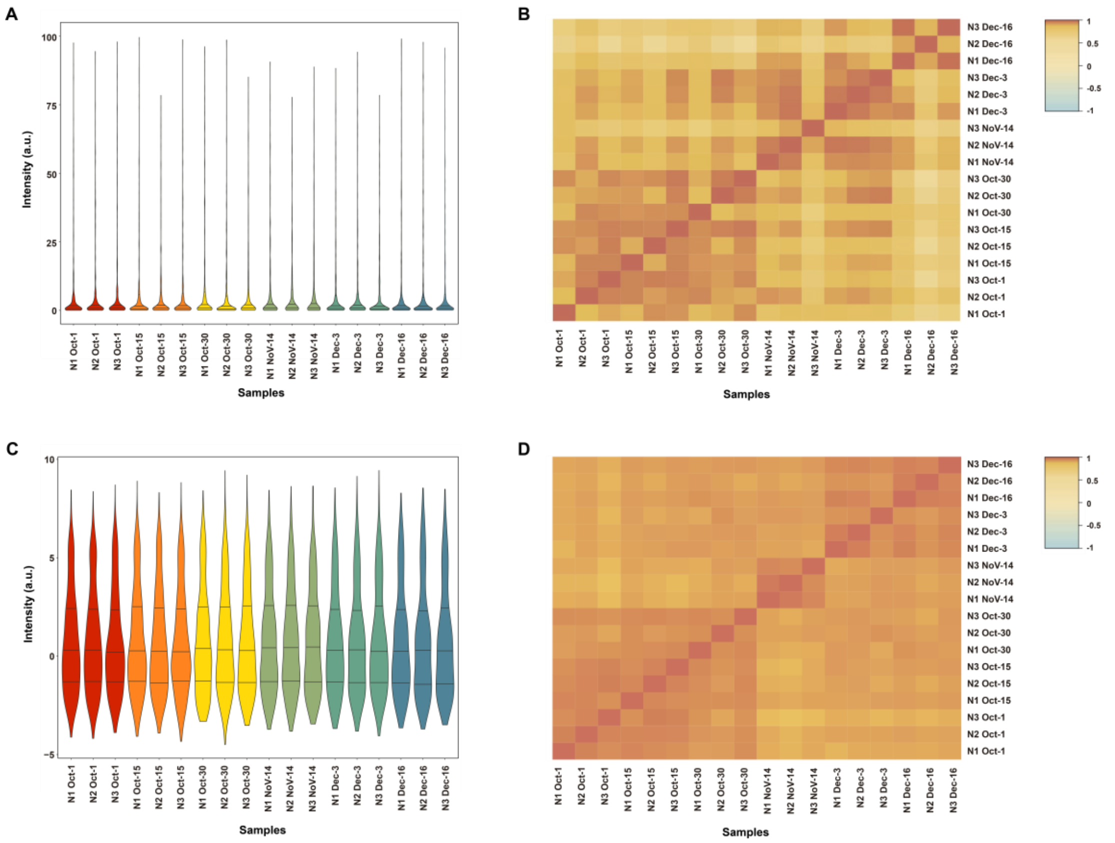


**Figure S1. Global distribution analysis of NMR-based metabolomic data**. **A**) Violin plots of raw data. Plots are colored according to the collection times (from October 1 to December 16, using a gradient from warm to cool colors). Horizontal lines inside each plot show the *upper* quartile, median, and *lower* quartile. N1: Node 1; N2: Node 2; N3: Node 3. **B**) Correlation heatmap of raw data. Sample labels are the same as in (A). The color key gradient represents the correlation coefficient from a strong negative correlation (−1.0, in blue) to a very strong positive correlation (1.0, in red). **C**) Violin plots of normalized data. Color key and sample labels are identical from those in Figure A. **D**) Correlation heatmap of normalized data. Sample labels and color key are identical from those in Figure B.

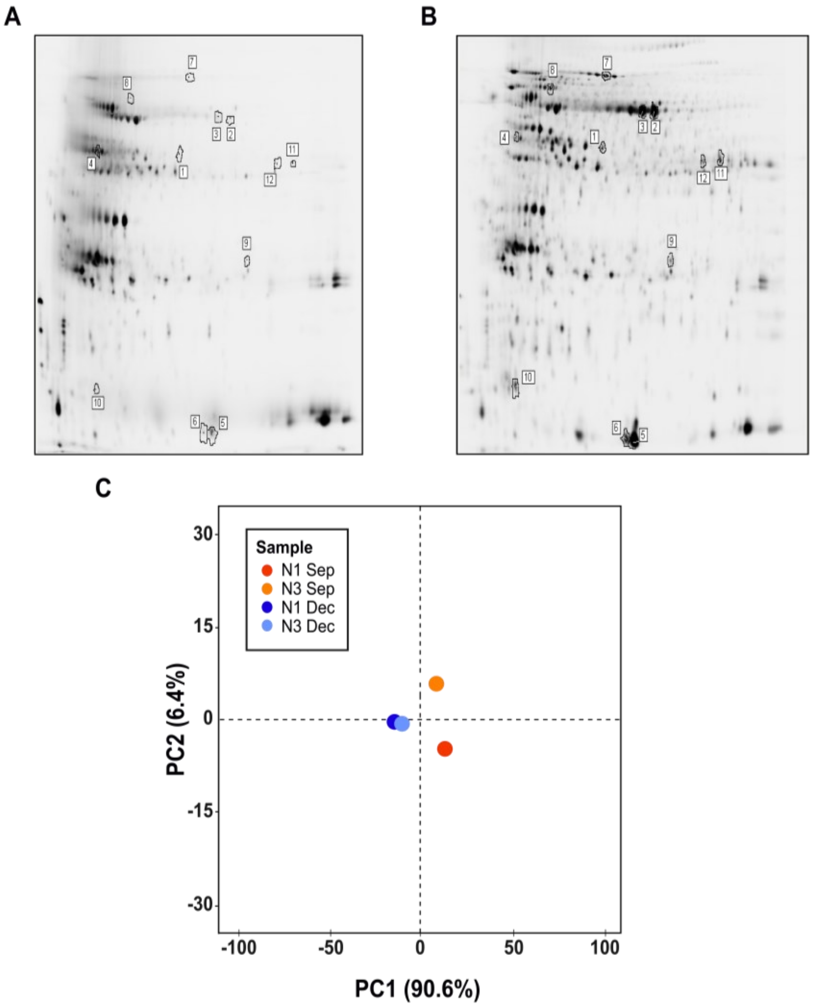


**Figure S2. Protein profile of strawberry leaves from different nodes collected on mid-September and mid-December. A)** Acrylamide gel showing the protein profile of a sample from Node 1 collected in September. Spots further identified by MS are labeled with numbers from 1 to 12. **B)** Acrylamide gel showing the protein profile of a sample from Node 1 collected in December. Spot labeling is identical to Figure A. **C)** Principal Component Analysis (PCA) of proteomic data from strawberry leaves grouped by node and time of collection. Nodes 1 and 3 are represented. Sample points are represented as the mean of three independent replicates performed of each node, and are labeled and colored according to the type of node and time of collection (Node 1 from September in red; Node 3 from September in orange; Node 1 from December in dark blue; Node 3 from December in light blue). PC1 or x-axis separates samples according to time of collection (mean of samples collected on December appear in negative values of the PC1, and mean of samples collected on September appear in positive values of the PC1). PC2 or y-axis separates samples collected on September (mean of samples from Node 1 appear in negative values of the PC2, and mean of samples from Node 3 appear in positive values of the PC2). Samples collected on December are scarcely influenced by PC2.
